# Interaction between static visual cues and force-feedback on the perception of mass of virtual objects

**DOI:** 10.1101/337899

**Authors:** Wenyan Bi, Jonathan Newport, Bei Xiao

## Abstract

We use force-feedback device and a game engine to measure the effects of material appearance on the perception of mass of virtual objects. We discover that the perceived mass is mainly determined by the ground-truth mass output by the force-feedback device. Different from the classic Material Weight Illusion (MWI), however, heavy-looking objects (e.g. steel) are consistently rated heavier than light-looking ones (e.g. fabric) with the same ground-truth mass. Analysis of the initial accelerated velocity of the movement trajectories of the virtual probe shows greater acceleration for materials with heavier rated mass. This effect is diminished when the participants lift the object for the second time, meaning that the influence of visual appearance disappears in the movement trajectories once it is calibrated by the force-feedback. We also show how the material categories are affected by both the visual appearance and the weight of the object. We conclude that visual appearance has a significant interaction with haptic force-feedback on the perception of mass and also affects the kinematics of how participants manipulate the object.

**CCS CONCEPTS:** • **Human-centered computing → Empirical studies in HCI; Empirical studies in interaction design**; *Empirical studies in visualization*;

**ACM Reference Format:** Wenyan Bi, Jonathan Newport, and Bei Xiao. 2018. Interaction between static visual cues and force-feedback on the perception of mass of virtual objects. In *Proceedings of*. ACM, New York, NY, USA, 5 pages.

## 1 INTRODUCTION

The most important function of material perception is to inform action. Past research on material perception has focused on a single modality (e.g., vision) with participants passively viewing images/videos and providing verbal reports. In everyday life, when we manipulate the objects, we often rely on both vision and touch to estimate their material properties. For example, when we reach out to pick a drinking bottle, both of its appearance and the tactile sensations (e.g., heaviness) will affect our judgment of its material properties. After we pick up the bottle, however, what we feel will rapidly update our belief in its material properties (perhaps it is made of plastic instead of glass?). It remains unclear how the visual information and tactile sensations interact to influence material perception. Recent advances in virtual reality (VR) and haptic device allow independent manipulations of visual and haptic inputs. Previous research has adapted such approach to understanding multi-modal sensory cue integration [Atkins et al. 2001; Scarfe and Glennerster 2015]. Relatively little has been explored in using the virtual environment to measure multimodal material perception.

In this paper, we aim to understand how visual appearance of the object affects its perceived mass and to test whether the classic Material Weight Illusion exists in an interactive virtual environment with force-feedback. In addition, we contribute a novel experimental paradigm to measure the perception of mass in an interactive virtual environment with realistic haptic force-feedback and high-fidelity visual rendering.

## 2 RELATED WORK

### 2.1 Mass perception of real and virtual objects

Existing work has found that several factors, such as size, depth cues, friction, and material properties, influence the perception of mass (see [Kawai et al. 2012] for a thorough review). The size has a significant effect on perceived mass [Buckingham and Goodale 2010; Mon-Williams and Murray 2000; Murray et al. 1999]. This is hypothesized by the expectation that a bigger object tends to be heavier. Such illusion is also experienced in relation to the objects’ material properties [Ellis and Lederman 1999]. Using the classic ‘Material Weight Illusion (MWI)’, Buckingham et al. showed that the visual material property affects the verbal judgements such that the heavier-looking object is rated to be lighter than its weight and lighter-looking object is rated to be heavier. More importantly, it also affects how much force participants initially apply to lift up an object such that they apply more force to lift up a heavier-looking object than a lighter-looking one even though they weigh the same [Buckingham et al. 2009]. However, after a few trials, the force is scaled back to the actual weight of the object, whereas the perceptual illusion remain [Flanagan and Beltzner 2000]. Our task is similar to that of the MWI where we independently manipulate the visual appearance of the object and its weight. But the crucial difference is that in our interactive virtual environment, the participants feel the gravity of the object before they can apply the force (see Discussion). So there will be no chance to scale the applied forces to the expected weight. Hence, if the hypothesis of the MWI is correct [Buckingham et al. 2009], we expect no illusion in our experiment. Another thread of previous research has shown that visually changing the physical behavior of the manipulated objects can create the haptic sensation in virtual environments ([Dominjon et al. 2005; Issartel et al. 2015]). To limit the effect of physical dynamics on mass perception but focus on material appearance, in this experiment we prevent the objects from bouncing when it collides with the surface.

### 2.2 Haptic and visual material perception

Research on material perception has focused on measuring properties such as surface gloss, roughness, translucency, viscosity, stiffness[Bi et al. 2018; Bi and Xiao 2016; Fleming et al. 2003; Guest and Spence 2003; Motoyoshi et al. 2007; Van Assen et al. 2018; Xiao et al. 2014], affective properties such as ‘fragility’ and ‘naturalness’ [Fleming et al. 2013], as well as material categories [Sharan et al. 2014] from visual inputs. Little is known about how visual and haptic information interact when we estimate material properties. Most of the work investigating the influence of haptic-visual interaction on material properties has focused on roughness [Guest and Spence 2003; Jones and O’Neil 1985; Lederman et al. 1986; Tiest and Kappers 2007] and compliance [Cellini et al. 2013; Kuschel et al. 2010; Massimiliano 2011; Massimiliano et al. 2011]. A few studies investigate whether haptic and visual representation of material properties and categories are similar [Baumgartner et al. 2013; Gaißert et al. 2010; Lederman et al. 1986; Overvliet and Soto-Faraco 2011; Tiest and Kappers 2007; Whitaker et al. 2008]. Paulun et al. investigated the effect of material properties on precision grip kinematics and found that materials affect the choice of local grasp points and the duration of the movement [Paulun et al. 2016]. Using a force-feedback device, Adams et al. examined whether human can integrate visual gloss with haptic ‘rubberiness’ when judging material properties [Adams et al. 2016]. But participants in their study do not have the chance to manipulate the objects while performing the task. By contrast, participants in our study can manipulate the object such as bringing it closer, picking it up, and dropping it.

## 3 EXPERIMENT

Eleven participants participated in the study. Using a phantom force-feedback device linked to a gaming environment, participants picked a virtual object from one place and dropped it to another. Then they were asked to rate the perceived mass of the object by comparing it to two reference cubes. In each trial, the object had a specific visual appearance and mass.

### 3.1 Apparatus and Experimental Setup

Figure 1A and Figure 1B show the apparatus used in this experiment. Our experimental set-up allowed concurrent and spatially aligned visual-haptic stimulus presentation. A monitor (ASUS VG248 LCD 24 inch) displaying the virtual scene was placed above the participant. The virtual scene was then reflected by a half-silvered mirror. A phantom force-feedback equipment (Geomagic Touch X, 3D systems) was set underneath the half-silvered mirror to provide the force feedback. During the experiment, the participant viewed the reflection of the virtual object through the half-silvered mirror, and they manipulated the object using a virtual probe in the scene which was spatially aligned with the stylus of the phantom. The head of the participant was fixed on a chin rest.

**Figure 1:**
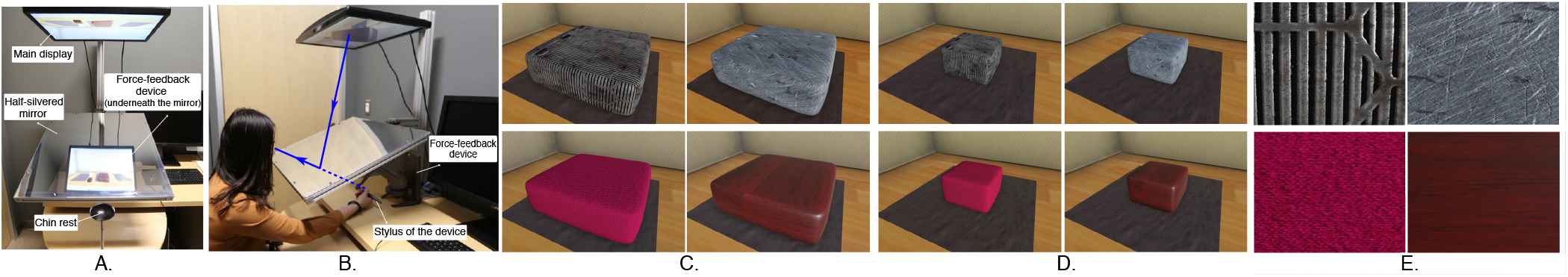
We create an interactive virtual environment which allows concurrent and spatially aligned visual-haptic presentation of virtual objects (A and B). Participants move the stylus of the force-feedback device (B) to manipulate the virtual object and feel its weight. The object is rendered with four materials (E: steel, stone, fabric, and wood) and two sizes (C and D). We independently vary the visual appearance (i.e. material and size) and the weight of the object, and measure how visual and haptic information affect the mass perception in the interactive virtual environment.

### 3.2 Stimuli

The virtual scene was rendered in Unity 3D game engine (2017.3.0f3). The scene was illuminated by a directional light and a reflection probe (see Figure 2). The game object was modeled as a rectangular prism with round corner and was rendered with four different surface materials: steel, stone, fabric, and wood (see Figure 1E) for a zoom-in view of the materials) using the Bitmap2Material asset for Unity. For each material type, the object was rendered into two sizes: a large size with the scale value [1.6, 0.4, 1.6] (Figure 1C)and a small size with a scale value [0.8, 0.4, 0.8] (Figure 1D). The force-feedback was rendered using Unity 5 Haptic Plugin for Geomagic OpenHaptics 3.3. There are several parameters provided in the plugin to control the haptic properties, all ranging from 0 to 1. In this experiment, we set the object stiffness to 1 and the damping value to 0.125. Both static and dynamic frictions were set to 1. We used three different mass levels (0.1, 0.5, 0.9) to simulate different levels of force-feedback. Higher mass level corresponded to larger force that was generated by the phantom.

**Figure 2:**
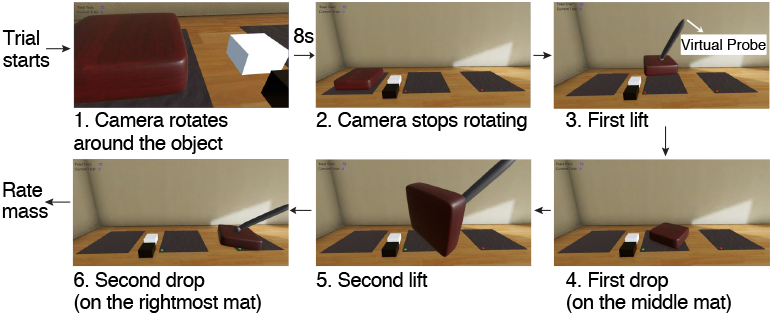
Task and procedure. In each trial, participants were first asked to watch the scene carefully while a camera rotating around the virtual object (1). Right after the camera stopped rotating (2), participants transported the object from the leftmost mat to the middle mat (3-4), then from the middle mat to the rightmost mat (5-6). During the transportation process, they could move the object closer to better observe its appearance.

### 3.3 Task

The experimental interface was created with Unity C# API, and the procedure is shown in Figure 2. In each trial, participants first saw a camera rotating around the virtual object. The rotation lasted for 9 seconds to allow participants better observe the material of the virtual object from multiple views. After the camera stopped rotating, participants started manipulating the object. They were instructed to move the stylus to first touch the upper surface of the The perception of mass of virtual objects object, and then press and hold a button to lift it. They wouldn’t feel any force feedback until the virtual probe touched the object and the button was pressed. After the manipulation, participants rated the mass of the object (0-100) that they just manipulated. In addition to the virtual object, the scene contained another two reference cubes: a white cube with a mass rating value of 0 and a black cube with a mass rating value of 100. Participants compared the mass of the virtual object to that of the two reference cubes to make an accurate rating. Each participant finished 72 trials in a random sequence (3 mass × 2 sizes× 4 materials × 3 repetitions).

## 4 RESULTS

### 4.1 Material appearance biases mass rating

To minimize the impact of individual range of perceptual scale between participants, the ratings are first normalized within each participant by linearly interpolating between 0 and 1 and then averaged across all participants. Figure 3 (C and D) plots the relationship between the physical and the perceived mass for each material type. First, perceived mass increases as the physical mass increases, indicating the weight of the object significantly affects perceived mass. Second, across all mass levels and scale sizes, the steel and stone are rated heavier than the fabric and wood material. A three way within-subject ANOVA with physical mass value, size, and material as factors reveals significant main effect of mass (*F*(2, 64) = 870.64, *p* < 0.001, MSE = 0.038, *η*^2^ = 0.981). Post-hoc comparisons suggest that objects with mass value 0.9 (*M* = 0.83, *SD* = 0.16) is rated significantly higher than that with mass value 0.5 (*M* = 0.43, *SD* = 0.16), which is also rated higher than that of 0.1 (*M* = 0.13, *SD* = 0.13)(*ps* < 0.05). Material also has amain effect (*F*(3, 96) = 16.32, *p* < 0.001, MSE = 0.307, *η*^2^ = 0.545), such that steel (*M* = 0.51, *SD* = 0.33) and stone (*M* = 0.48, *SD* = 0.32) are consistently rated significantly heavier than fabric (*M* = 0.42, *SD* = 0.41) and wood (*M* = 0.44, *SD* = 0.32) (*ps* < 0.05). No other significance is revealed.

**Figure 3:**
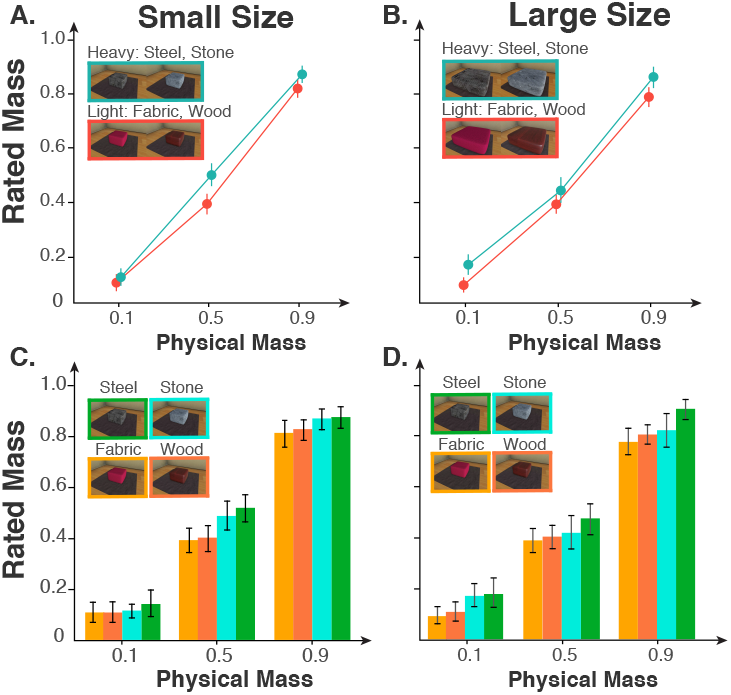
Rating of perceived mass. A. Perceived mass rating versus physical mass for two material categories of the small objects. Cyan colored line represents heavier-looking materials and orange colored line represents lighter-looking materials. B. Same for the large objects. C. Perceived mass rating for four materials (steel, stone, fabric, wood) of the small objects. D. Same for the large objects.

We then group the four materials to ‘heavier-looking’ (stone and steel) and ‘lighter-looking’ (fabric and wood) (Figure 3, A and B). The cyan colored line is consistently above the orange colored line, suggesting heavier-looking object is perceived heavier for each mass value. Again, physical mass value and material type both show significant main effects (*ps* < 0.001). There is also a significant interaction between size and physical mass value (F(2, 130) = 382.02, MSE = 0.014, *p* < 0.001, *η*^2^ = 0.855) such that bigger objects (*M* = 0.14, *SD* = 0.14) are rated as heavier than smaller objects (*M* = 0.11, *SD* = 0.12) when the physical mass is low (mass = 0.1). But this difference disappears for heavier objects (mass > 0.1).

These results show that mass perception is mainly affected by the haptic information. But visual appearance also has an effect such that heavier-looking objects are perceived as heavier than lighter-looking objects.

### 4.2 Visual appearance affects manual activity

It is possible that the expected weight of the object (which might be induced by the visual appearance) influences how much force participants prepare to apply to lift the object. We can infer this information by computing the initial accelerated velocity of the trajectory of the phantom stylus. We hypothesize that the accelerated velocity does not change within a very short time window (*δt* = 0.01 *seconds*). Therefore, we can use Newton’s second law of movement to calculate the accelerated velocity at the moment when the participate lift up the object. Figure 4 compares the accelerated velocities within the initial 0.1seconds for the heavy-looking and light-looking materials. The results show that the accelerated velocity for heavy-looking materials is higher than that of the light-looking one during the first lift. But this effect disappears when the participants lift the object the second time.

**Figure 4:**
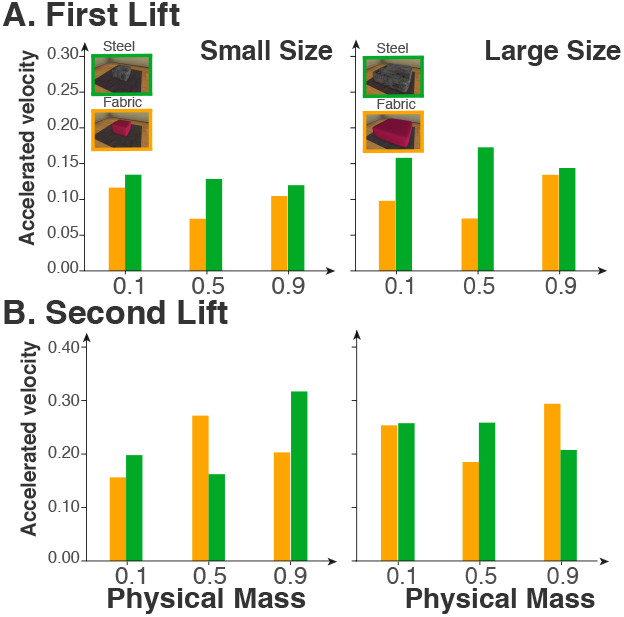
The comparison of initial accelerated velocities of the heavy-looking (steel, coloredingreen) andlight-looking (fabric, colored in orange) objects. A. First lift. Across all three mass levels, the accelerated velocity of the steel is larger than that of the fabric. B. No systematical differences in the second lift.

### 4.3 Material categorization is affected by both force-back and appearance

Figure 5 plots the frequency of the reported material names across all participants and all scale sizes as a word cloud. First, the figure shows that material categories on average remain the same while the physical mass increases. It also shows that within each mass/visual condition, the material name reported indicates the perceived mass. For example, the participants who provide the name ‘cloth cushion’ for the stone object also give lower mass rating than those who provide the name ‘stone’. This shows that the material category is consistent with the perceived mass in general. Together, this demonstrates that material category is predominately determined by visual appearance. But the physical mass of the object also has a small effect. For example, when the steel object feels heavy, participants consistently name it ‘metal’ or ‘steel’. But when it feels lighter, the names include ‘aluminum’ and ‘copper’.

**Figure 5:**
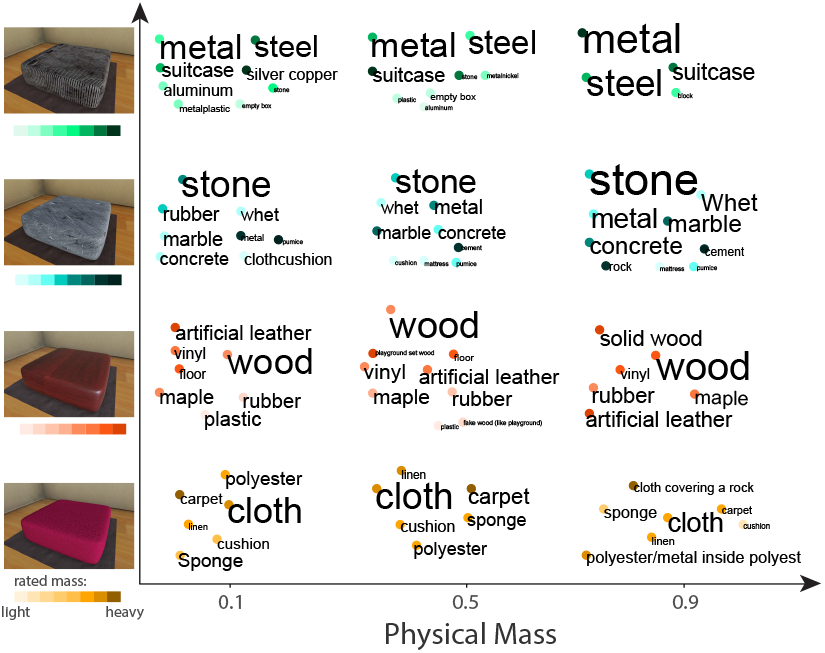
Word cloud based categorization analysis. X-axis represents the physical mass value, and Y-axis refers to material categories. The word cloud is summarized across all participants and size values. Material names with bigger fonts have higher frequency, and the differences between the font sizes indicate the true differences in the frequency. The colored dots underneath each material name refer to the mean perceived mass across all participants that provided this name. Darker color refers to higher perceived mass, and the saturation indicates the relative ranking.

## 5 DISCUSSION

### 5.0.1 Relationship with classic material weight illusion (MWI)

This paper finds that heavier-looking object is rated to be heavier, which is opposite from the MWI using real objects. This might due to the time course of how the participants lift the virtual objects. When lifting a real object, participants can prepare the gripping force before touching, so the gravity on the object and the applied force concurrently take effect as soon as the object is lifted up. Thus, the applied force has the opportunity to be scaled up to the expected weight. In our experiment, in order to lift the virtual object, the participants first need to press the button on the phantom stylus (see Figure 1B) when the corresponding virtual probe in the virtual scene (see Figure 2(3)) touches the object. Due to this action, it is impossible to prepare the lifting force while holding the object to prevent it from falling off. As soon as the button is pressed, the gravity of the object takes effect. Thus, the participants have to passively react to the gravity instead of actively applying a lifting force. Future study should consider this effect when using haptic device to study mass perception. Another possibility is that some of our objects look hollow (steel), which might induce a belief that the objects have an uneven density and thus affect the results.

### 5.0.2 Force feedback calibration on verbal judgements and actions

We also find that the visual appearance affects the lifting actions for the first lift, but not for the second. One possibility is that the action has been calibrated by the first lift and thus becomes more accurate in the second lift. However, the verbal rating is still affected by the visual appearance even after calibration. This is consistent with previous findings showing that manual activities could be accurately calibrated while verbal ratings are under influence by cognitive factors [Buckingham and Goodale 2010; Li et al. 2011]. Future studies should include both verbal reporting and manual activity to measure material perception.

### 5.0.3 Interaction between size and mass

We find a reversed size-weight illusion only when the ground-truth mass value is low. One explanation is that according to Weber’s law, the perceptual scale is logarithmic. Therefore, participants might be more sensitive to objects with low mass value, but when the mass becomes larger the discrimination threshold of the difference is higher.

## 6 CONCLUSION

Using a haptic force-feedback device in the virtual environment, we find that visual material properties have a significant effect on perceived mass. Different from the material weight illusion, the participants consistently rate heavier-looking objects to be heavier. Analysis of the initial accelerated velocity of the virtual probe shows higher acceleration for materials with higher perceived mass than those with lighter perceived mass. This effect is diminished when the participants pick up the object for the second time. This work provides new insights of using haptic device and virtual environments to study perception of physical properties of objects.

## ACKNOWLEDGMENTS

The authors would like to thank George Gu and Erik Daniel Whipp for helpful discussions about the set-up, task, and the softwares. The perception of mass of virtual objects

